# MtMOT1.2 is responsible for molybdate supply to *Medicago truncatula* nodules

**DOI:** 10.1101/272112

**Authors:** Patricia Gil-Díez, Manuel Tejada-Jiménez, Javier León-Mediavilla, Jiangqi Wen, Kirankumar S. Mysore, Juan Imperial, Manuel Gonzalez-Guerrero

**Author notes:** **One Sentence summary:** MtMOT1.2 mediates molybdate transfer from the vasculature to nitrogen-fixing nodules. **List of author contributions:** P.G-D. carried out most of the experiments. Yeast transport assays were performed by M.T-J. J. L-M. studied the complementation with molybdate of the **mot1.2-1** mutant. J.W. and K.S.M performed M. truncatula mutant screening and isolated the **mot1.2-1** allele. M.T-J, J.I., and M.G-G. designed the experiments, analysed the data, and wrote the manuscript with input from all the other authors. Corresponding author: Manuel González-Guerrero, Manuel Tejada-Jiménez.

## Abstract

Symbiotic nitrogen fixation in legume root nodules requires a steady supply of molybdenum for synthesis of the iron-molybdenum cofactor of nitrogenase. This nutrient has to be provided by the host plant from the soil, crossing several symplastically disconnected compartments through molybdate transporters, including members of the MOT1 family. MtMOT1.2 is a *Medicago truncatula* MOT1 family member located in the endodermal cells in roots and nodules. Immunolocalization of a tagged MtMOT1.2 indicates that it is associated to the plasma membrane and to intracellular membrane systems, where it would be transporting molybdate towards the cytosol, as indicated in yeast transport assays. A loss-of-function *mot1.2-1* mutant showed reduced growth compared to wild-type plants when nitrogen fixation was required, but not when nitrogen was provided as nitrate. While no effect on molybdenum-dependent nitrate reductase activity was observed, nitrogenase activity was severely affected, explaining the observed difference of growth depending on nitrogen source. This phenotype was the result of molybdate not reaching the nitrogen-fixing nodules, since genetic complementation with a wild-type *MtMOT1.2* gene or molybdate-fortification of the nutrient solution, both restored wild-type levels of growth and nitrogenase activity. These results support a model in which MtMOT1.2 would mediate molybdate delivery by the vasculature into the nodules.

## INTRODUCTION

Symbiotic nitrogen fixation carried out by the legume-rhizobia partnership is one of the main sources of assimilable nitrogen in natural ecosystems and in sustainable agriculture (Downie, 2014). The symbiosis is established in differentiated root organs, the nodules, developed after a complex chemical exchange between the symbionts (Oldroyd, 2013). Nodule inner cells are infected by rhizobia in an endocytic-like process that results in organelle-like structures, the symbiosomes (Vasse, 1990; Limpens et al., 2009). There, surrounded by a plasmalemma-derived membrane, the symbiosome membrane, rhizobia will differentiate into nitrogen-fixing bacteroids (Roth, 1989, Vasse, 1990). As a result, dedicated membrane transporters would be required to transfer the fixed nitrogen to the host plant, while the bacteroid receives photosynthates and mineral nutrients (Udvardi and Poole, 2013).

Transition metal nutrients, such as iron (Fe), copper (Cu), zinc (Zn), or molybdenum (Mo), are required in relatively large amounts for symbiotic nitrogen fixation (O’Hara, 2001; Brear et al., 2013; González-Guerrero et al., 2014). While Fe, Cu, and Zn are employed as cofactors of multiple enzymes present in the nodule (Brear et al., 2013; González-Guerrero et al, 2014), Mo is required by just two when nitrogen fixation is active. One of them is a xanthine dehydrogenase that might be required for nitrogen delivery out of the nodules (Kaiser et al., 2005); while the other is nitrogenase, the protein complex directly responsible for converting N_2_ into NH_4_^+^ by the bacteroids. In this enzyme, Mo is a key element of its unique Fe-Mo cofactor (FeMoco) (Rubio and Ludden, 2005). Consequently, Mo uptake and delivery to the nodules is an essential process for legumes and for symbiotic nitrogen fixation.

Mo is present in soil as molybdate, a close structural analogue to sulfate (Stiefel, 2002). In fact, for many years it was believed that molybdate was mainly transported by sulfate carriers, since, under the right conditions, they can also carry molybdate across membranes (Stout et al., 1951; Mendel and Hansch, 2002; Kaiser et al., 2005). However, more recently, molybdate-specific transporters have been identified (Tejada-Jiménez et al., 2007; Tomatsu et al., 2007; Tejada-Jiménez et al., 2011). Among them, the better known is the MOT1 family of transporters (Tejada-Jimenez et al., 2007; Tomatsu et al., 2007; Baxter et al., 2008). These proteins are evolutionarily related to sulfate transporters of the SULTR family (Tejada-Jimenez et al., 2007). However, the conserved STAS domain common to the latter proteins is not present in the MOT1 family members, which are characterized by the domains: P-PVQPMKXIXA-A and FG-MP-CHGAGGLA-QY-FGGR-G (Tejada-Jimenez et al., 2007). In Arabidopsis, two *MOT1* genes have been found: *AtMOT1*, involved in Mo uptake (Tomatsu et al., 2007; Baxter et al., 2008); and *AtMOT2*, associated to the vacuole, and playing a role in inter-organ molybdate allocation (Gasber et al., 2011).

On an average, legume genomes encode more copies of MOT1 gene family members than other dicots. This could be the result of evolutionary pressures to expand this family to account for the increased demand of Mo in symbiotic nitrogen fixation (O’Hara et al., 2001). For instance, *Glycine max* has seven members; *Phaseolus vulgaris*, four; and *Medicago truncatula* has five (MtMOT1.1 to MtMOT1.5). Two legume MOT1 proteins have been characterized to date (Gao et al., 2016; Duan et al., 2017; Tejada-Jimenez et al., 2017). LjMOT1 would be responsible for molybdate uptake from soil and its distribution to plant sink organs (Gao et al., 2016; Duan et al., 2017), in a role similar to that played by AtMOT1 (Tomatsu et al., 2007). This is indicated by the expression of this transporter in the epidermis and in the root vasculature, as well as by the reduction of Mo content in nodules and leaves in mutant plants (Duan et al., 2017). However, LjMOT1 would not primarily be involved in molybdate delivery to the nodules, since lack of this transporter has no significant effect on nitrogen fixation capabilities. In contrast, mutation of the other characterized legume MOT1 protein, nodule-specific MtMOT1.3, results in nearly a total loss of nitrogenase activity (Tejada-Jiménez et al., 2017). Immunolocalization of this transporter in the plasma membrane and characterization of its transport capabilities indicate that its main function would be introducing molybdate into nodule cells. However, *mot1.3-1* mutant nodules still accumulate more Mo than wild-type ones, suggesting that the delivery of this metal to the nodules is not impaired by *MtMOT1.3* mutation (Tejada-Jiménez et al., 2017). Consequently, we still need to identify the transporters responsible for molybdate release from the vasculature into the nodules. This function would require two types of transporters: a MOT1-like for uptake from the vessels, and a yet-to-be-defined molybdate efflux protein (perhaps a sulfate transporter) to extrude Mo into the nodule apoplast.

Among the four remaining MOT1 transporters in *M. truncatula*, MtMOT1.1 is the most closely related to LjMOT1, and would likely play a similar role (Tejada-Jiménez et al., 2017). *MtMOT1.4* and *MtMOT1.5* are expressed all over the plant, while *MtMOT1.2* is the only one of these four genes that is expressed exclusively in roots and nodules, with a maximum of expression in the latter (Tejada-Jimenez et al., 2017). Here, we characterize the function of MtMOT1.2 as a likely candidate for molybdate uptake from the vasculature by endodermal cells. MtMOT1.2 is located in the endodermis of nodule and root vascular cylinders, in the plasma membrane and in an endomembrane compartment. It shows molybdate uptake capabilities in yeast, and its mutation in *M. truncatula* leads to a reduction in nitrogenase activity in nodules, likely the result of the reduction in molybdate delivery to the nodules. Its function seems to be relevant for symbiotic nitrogen fixation, with no major role being played under non-symbiotic conditions, given how its mutation has no effect on plants grown on nitrate or on nitrate reductase activity. This work represents a further step towards understanding how molybdate in particular, and transition metals, in general, are delivered to legume nodules, and represents the first case in which a metal transporter has been associated with root-to-nodule vascular metal delivery.

## RESULTS

### MtMOT1.2 is a molybdate transporter

It has been reported that proteins belonging to MOT1 family are involved in the transport of the oxyanion molybdate into the cytosol of cells (Tejada-Jimenez et al., 2013). Members of MOT1 family showed a high sequence similarity to other molybdate transporters, including the signature motives of MOT1 family (Tejada-Jimenez et al., 2007). To confirm that MtMOT1.2 was able to transport molybdate, a yeast expression system was used, since these *Saccharomyces cerevisiae* is among the rare organisms lacking Mo-containing proteins and having no Mo transporters (Mendel and Bittner, 2006). When grown in the presence of molybdate, yeast expressing *MtMOT1.2* were able to accumulate Mo following a Michaelian kinetics (Fig. 1A), with a *V*_*max*_ of 155 ± 12 pmol 10^-6^ cells h^-1^ and a *k*_*1/2*_ of 488 ± 105 nM. This transport was not inhibited by the structural analogue sulfate, even at concentrations up to 4,000 times higher (Fig. 1B).

**Figure 1.**
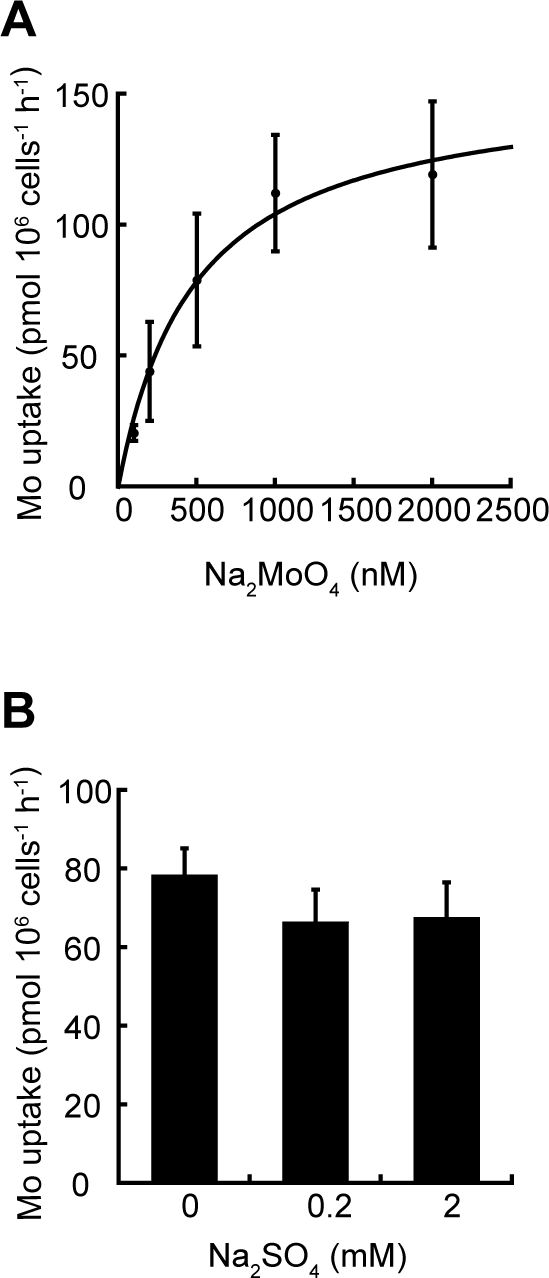
*Medicago truncatula* Molybdate Transporter 1.2 (MtMOT1.2) introduces molybdate towards the cytosol. (A) Molybdate uptake by *Saccharomyces cerevisiae* strain 31019b transformed with PDR196 vector containing *MtMOT1.2* coding sequence and grown at 28°C. Values for the pDR196 empty vector were substracted from the data. Data were fitted using Michaelis constant k_1/2_= 488 ± 105 nM and maximum speed v_max_= 155 ± 12 pmol 10^6^ cells^-1^ h^-1^. Data are the mean ± SD. (B) Effect of sulfate on molybdate uptake by *S. cerevisiae* strain 31019b transformed with PDR196 containing the *MtMOT1.2* coding sequence. Data sets consist of *S. cerevisiae* incubated with 500 nM Na_2_MoO_4_ and 0, 0.2, or 2 mM Na_2_SO_4_. Data are the mean ± SD.

### *MtMOT1.2* is located in the plasma membrane and intracellular compartments in the endodermal/periccle layer in roots and nodules

According to Symbimics database (https://iant.toulouse.inra.fr/) (Roux et al., 2014) and previously reported data (Tejada-Jiménez, 2017), *MtMOT1.2* is expressed in nodules and roots of *M. truncatula.* In order to identify the tissues in which its expression peaks, 1,446 bp upstream of the start codon of *MtMOT1.2* were fused to a β glucuronidase (*gus*) gene. *M. truncatula* plants were transformed with this genetic construct and GUS activity was visualized at 28 days-post-inoculation (dpi). The results confirmed that *MtMOT1.2* was expressed in nodules and roots (Fig. 2A). Root sections showed that most of the GUS activity was confined to the endodermal layer around the root vessels (Fig. 2B). Similarly, *MtMOT1.2* expression in the nodules was located around the nodule vasculature (Fig. 2C), and no expression was observed in the inner nodule regions, even when they were clarified with bleach (Supplemental Fig. S1).

**Figure 2.**
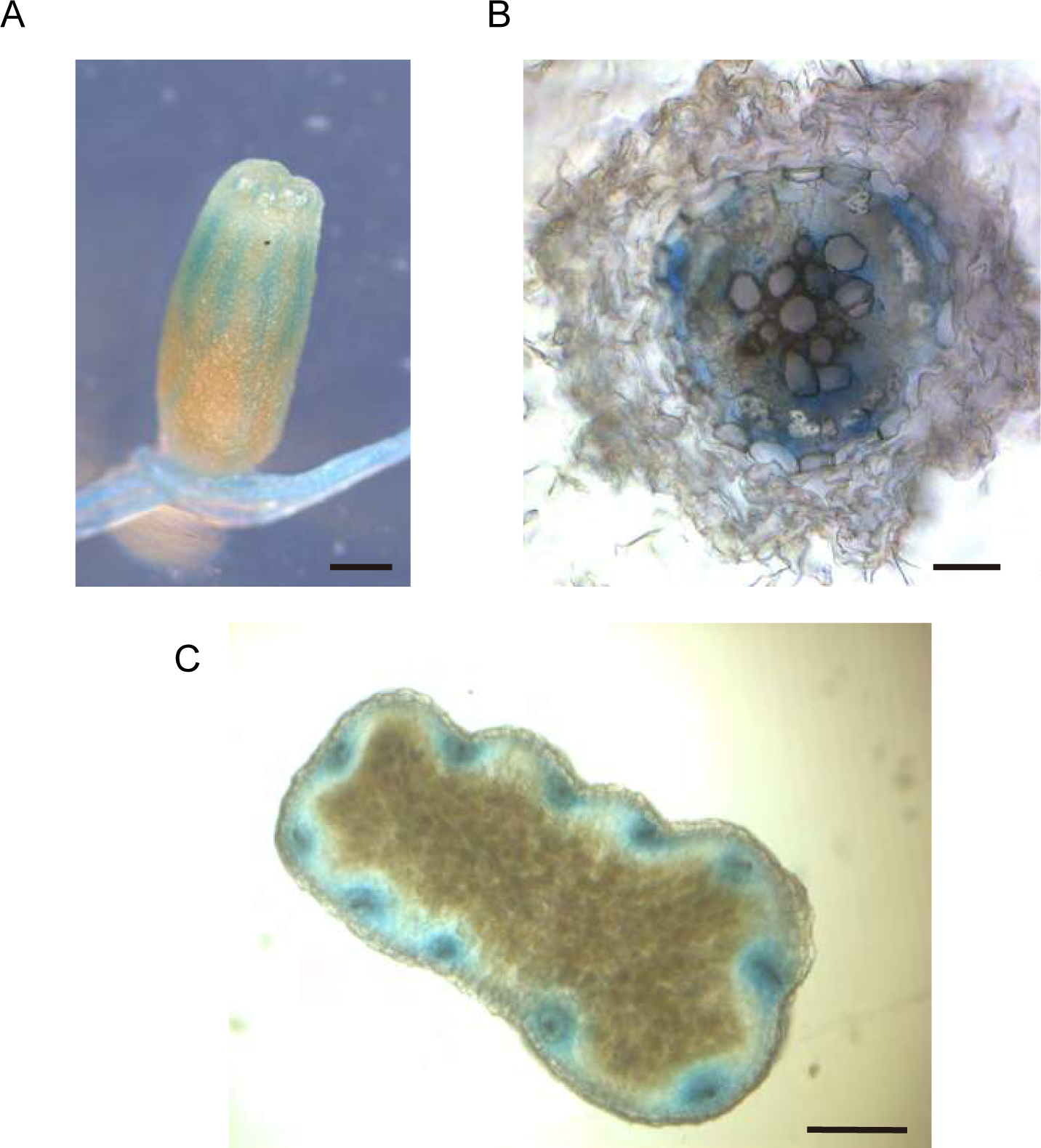
*Medicago truncatula Molybdate Transporter 1.2* (*MtMOT1.2*) gene is expressed in the around the vessels in roots and nodules. (A) β-glucuronidase (GUS) staining of *M. truncatula* roots and nodules from 28 days-post-inoculation (dpi) plants transiently expressing the *gus* gene under the control of *MtMOT1.2* promoter region. Scale bar = 0.5 mm. (B) GUS activity localization in a cross section of a 28 dpi root from *M. truncatula* plants transiently expressing the *gus* gene under the control of *MtMOT1.2* promoter region. Scale bar = 0.025 mm. (C) GUS activity localization in a cross section of a 28 dpi nodule from *M. truncatula* plants transiently expressing the *gus* gene under the control of *MtMOT1.2* promoter region. Scale bar = 25 µm.

Immunolocalization of MtMOT1.2 was performed by fusing the full genomic region (comprising from 1,446 bp upstream of the start codon to the last codon before the stop) to three hemagglutinin (HA) epitopes. Localization of the MtMOT1.2-HA protein was carried out with a mouse anti-HA primary antibody and a secondary anti-mouse Alexa594-conjugated antibody. The plants were inoculated with a *S. meliloti* strain constitutively expressing green fluorescent protein (GFP), and DNA was stained blue with 4’-6-diamino-phenylindole (DAPI). The result of this staining showed that MtMOT1.2-HA was located in a cell layer around the root and nodule vessels (Fig. 3A and B), thus validating the *gus*-reporter assays. The detection of autofluorescence bands corresponding to the Casparian strip suggests that these cells form the endodermis of the nodule vessels (Supplementary Fig. S2). The Alexa594 signal was observed in two locations within a cell: in the periphery of the cells and in a perinuclear region (Fig. 3A). In the root, MtMOT1.2-HA had a similar cellular distribution (Fig. 3B). This pattern of detection was not the result of autofluorescence detected in the Alexa594 emission channel, since negative controls with exactly the same conditions did not show any signal in this emission range (Supplemental Fig. S3).

**Figure 3.**
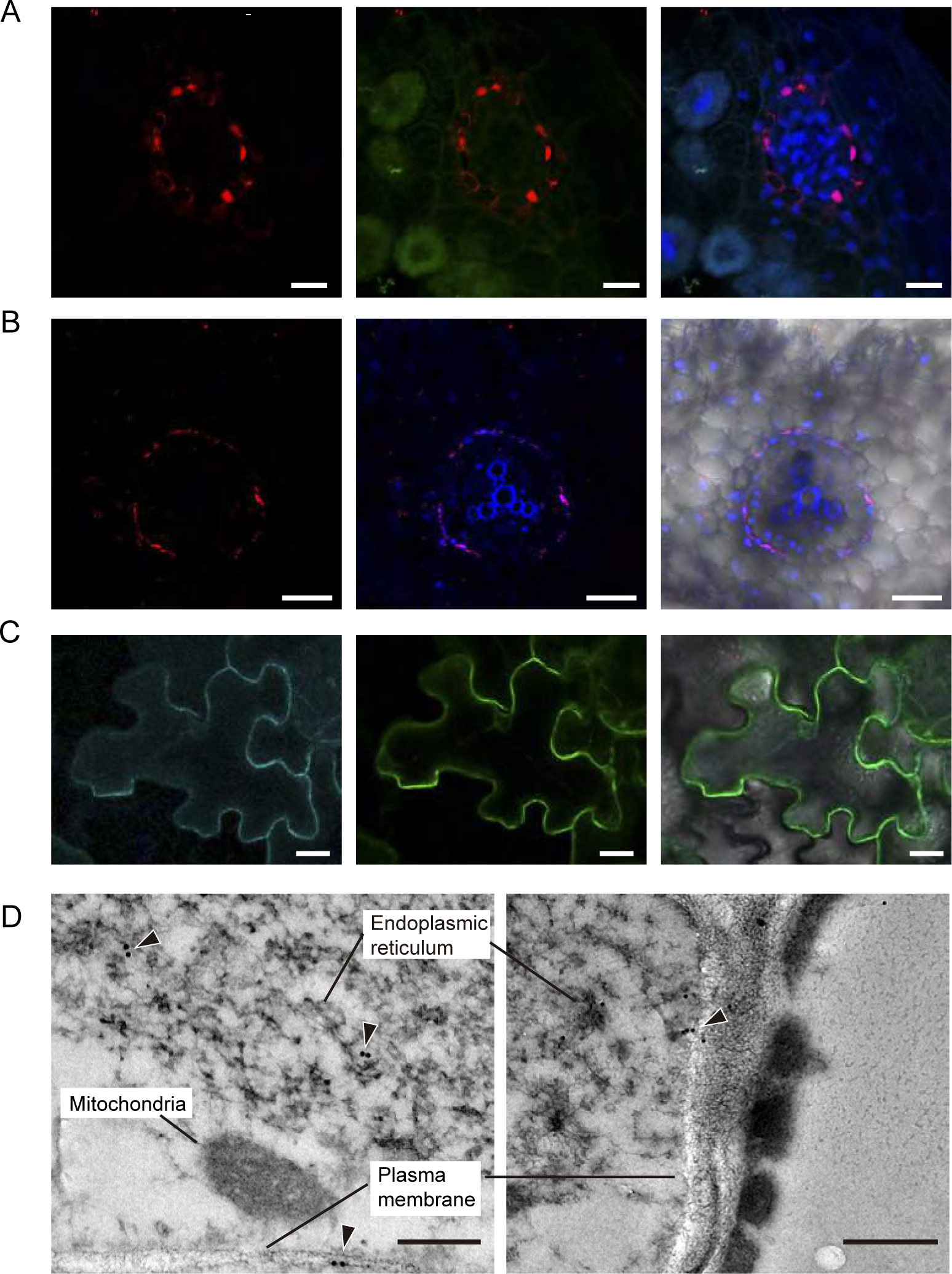
*Medicago truncatula* Molybdate Transporter 1.2 (MtMOT1.2) is located in the plasma membrane and an endomembrane compartment in endodermal cells. (A) Cross-section of a 28 days-post-inoculation (dpi) *M. truncatula* nodule expressing *MtMOT1.2-HA* and inoculated with a *Sinorhizobium meliloti* strain constitutively expressing the green fluorescent protein (GFP) (green). MtMOT1.2-HA was detected using an Alexa594-conjugated antibody (red). DNA was stained with DAPI (blue). Left panel, localization of MtMOT1.2-HA; central panel, overlay with the green (*S. meliloti)* channel; right panel, overlay of the central panel with the DAPI-stained DNA. Scale bars = 10µm. (B) *M. truncatula* root expressing *MtMOT1.2-HA*. MtMOT1.2-HA was detected using an Alexa594-conjugated antibody (red). DNA was stained with DAPI (blue). Left panel, localization of MtMOT1.2-HA; central panel, overlay with the DAPI-stained DNA and the xylem autoflorescence; right panel overlay with the transillumination image. Scale bars = 50 µm. (C) Transient co-expression of MtMOT1.3-GFP (green) and AtPIP2-CFP (cyan) in *Nicotiana benthamiana* leaves. Left panel, AtPIP2-CFP signal, middle panel MtMOT1.2-GFP signal; right panel overlay of the two channels with the transillumination image. Scale bars = 25 µm. (D) Subcellular localization of MtMOT1.2-HA in nodule vessels using a gold-conjugated anti-HA antibody. Arrowheads indicate the position of the gold particles. Scale bars = 250 nm and 500 nm.

To obtain further detail on the subcellular distribution of MtMOT1.2, *Nicotiana benthamiana* leaves were co-agroinfiltrated with a C-terminal GFP-tagged MtMOT1.2 and the plasma membrane-marker AtPIP2 labelled with cyan fluorescent protein (CFP). Figure 3C shows that both signals co-localize, indicating that in *N. benthamiana* MtMOT1.2 is located in the plasma membrane. Again, these signals were not due to autofluorescence, since neither GFP nor CFP were detected in leaves expressing just AtPIP2-CFP, or MtMOT1.2-GFP, respectively (Supplemental Fig. S4). MtMOT1.2-HA localization was also determined with transmission electron microscopy and a gold-conjugated secondary antibody (Fig. 3D). In these sections, the epitope was detected in the plasma membrane and in an intracellular membrane compartment, likely the endoplasmic reticulum. No gold particles were detected in control nodules (Supplemental Fig. S5).

### MtMOT1.2 is required for molybdate delivery to nitrogen fixing root nodules

A *M. truncatula Transposable Element from N. tabacum* (*Tnt1*) insertion lines (Tadege et al., 2008) were used for a reverse genetics screening (Cheng et al., 2011; Cheng et al., 2014) to identify a mutant line (NF9961, *mot1.2-1*) carrying *Tnt1* in *MtMOT1.*2 to determine the physiological role of MtMOT1.2. *Tnt-1* insertion in the mutant *mot1.2-1* is located in the second exon of the gene, 1,576 bp downstream of the start codon (Fig 4A). No *MtMOT1.2* transcript was detected by RT-PCR in either roots or nodules of *mot1.2-1* (Fig. 4B). Since *MtMOT1.2* was expressed in roots of plants inoculated and non-inoculated with *S. meliloti, mot1.2-1* phenotype was assessed under both conditions.

**Figure 4.**
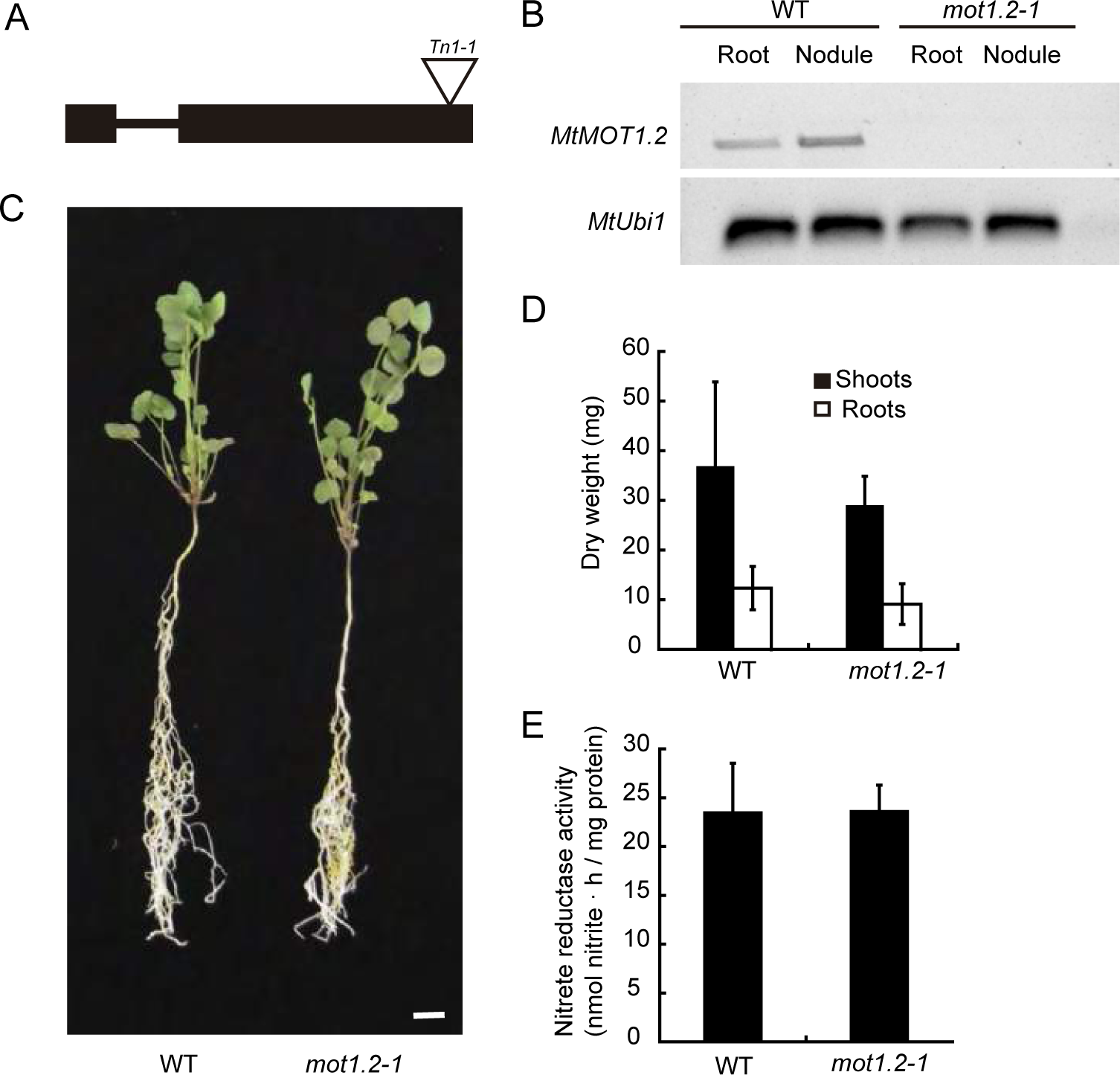
*Medicago truncatula* Molybdate Transporter 1.2 (MtMOT1.2) is not required for growth under non-symbiotic conditions. (A) Position of the *Transposable element from Nicotiana tabacum1* (*Tnt1*) insertion within the *MtMOT1.2* gene. (B) RT-PCR amplification of *MtMOT1.2* coding sequence in 28 days-post-inoculation roots and nodules of *M. truncatula* wild type (WT) or mutant (*mot1.2-1*) plants. *Ubiquitin carboxyl-terminal hydrolase1* (*MtUb1*) was used as a control. (C) Growth of representative WT and *mot1.2-1* plants watered with KNO_3_. Scale bar = 1 cm. (D) Dry weight of shoots and roots. Data are the mean ± SD of at least 6 independently transformed plants. (E) Nitrate reductase activity. Nitrate reduction was measured in duplicate. Data are the mean ± SD.

When the plants received an assimilable form of nitrogen (KNO_3_) in their nutrient solution, and no *S. meliloti* inoculum was added, no significant differences in growth (Fig. 4C) or in biomass production (Fig. 4D) were observed between wild-type and mutant plants. Since nitrate was the sole nitrogen source for these plants, they would require the activity of the Mo-containing enzyme nitrate reductase to grow (Bernard and Habash, 2009), and any deffect on Mo uptake or source-to-sink delivery in these plants would lead to a reduction of Mo-dependent enzymatic activities. However, nitrate reductase activity in *mot1.2-1* plants watered with KNO_3_ was equivalent to that of wild-type plants (Fig. 4E). No significant change was observed even when no molybdate was added to the nutrient solution (Supplemental Fig. S6).

In contrast, under symbiotic conditions, when the plant depends on symbiotic nitrogen fixation as the sole source of nitrogen, *mot1.2-1* plants showed reduced growth when compared to the controls (Fig. 5A), with smaller nodules (Fig. 5B) and a biomass reduction of 54% and 38% in shoot and root, respectively (Fig. 5C). The reduction of growth and nodule size did not seem to be the consequence of alterations on nodule development, or deffects in nodulation kinetics (neither the number of nodules per plant, nor the rate of nodulation were affected; Supplemental Fig. S7). Growth reduction in *mot1.2-1* plants is the likely result of a reduction of nitrogenase activity in those nodules, which was only 12% of that in wild-type plants (Fig. 5D). Mutant nodules exhibited a significant reduction in Mo content, while no significant changes were observed in roots (Fig. 5E). This phenotype was the result of the mutation of *MtMOT1.2*, since transforming *mot1.2-1* with *MtMOT1.2-HA* improved growth and restored wild-type levels of nitrogenase activity (Fig. 5). Similarly, increasing molybdate content in the nutrient solution restored wild-type growth and nitrogenase activity in *mot1.2-1* nodules (Supplemental Figure S8).

**Figure 5.**
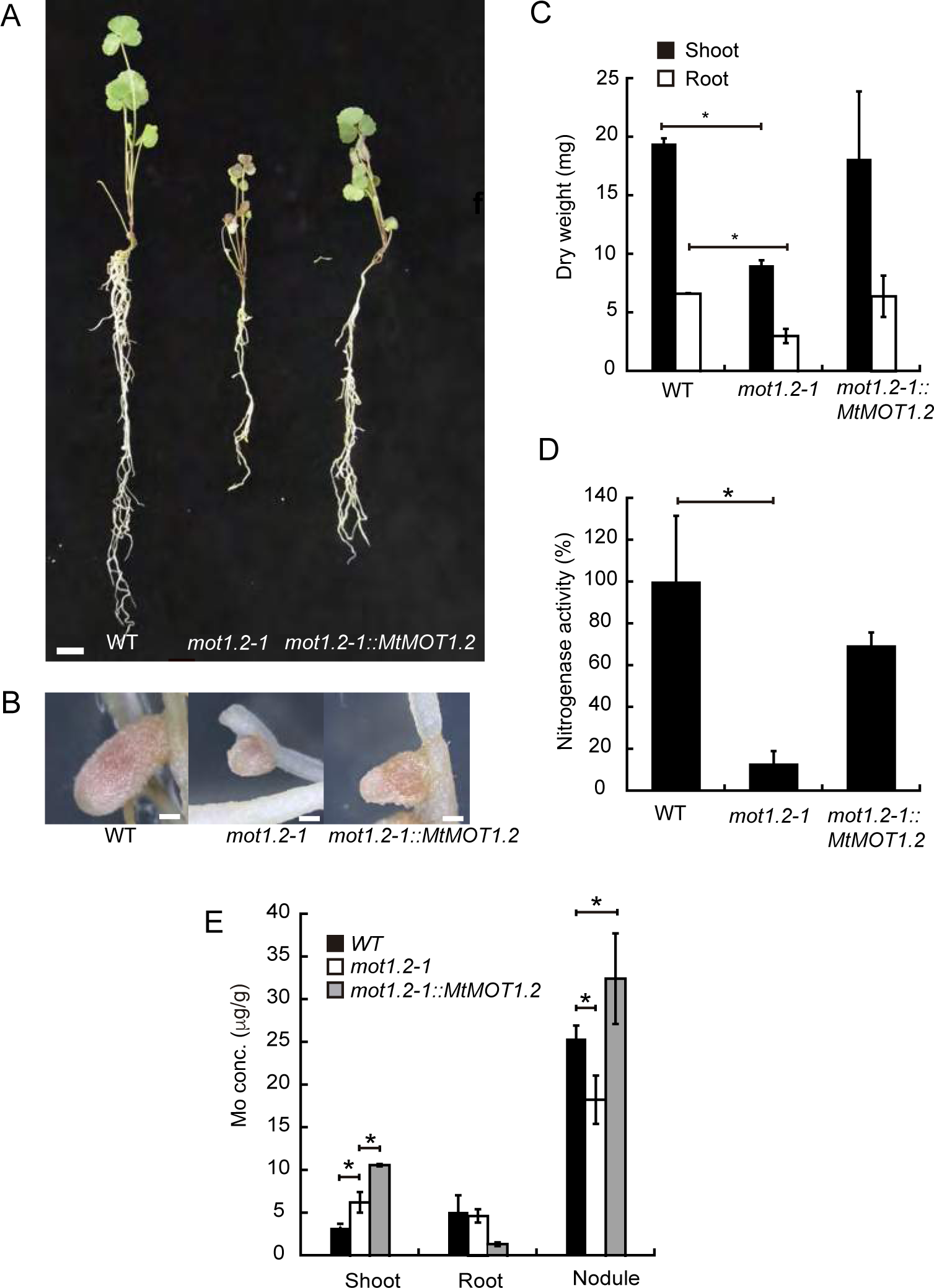
*Medicago truncatula* Molybdate Transporter 1.2 (MtMOT1.2) is required for symbiotic nitrogen fixation. (A) Growth of representative WT, *mot1.2-1*, and *mot1.2-1* transformed with *MtMOT1.2-HA* plants 28 days-post-inoculation (dpi). Scale bar = 1 cm.(B) Detail of representative 28 dpi nodules from WT, *mot1.2-1*, and *mot1.2-1* transformed with *MtMOT1.2-HA* plants. Scale bars = 50 mm. (C) Dry weight of shoots, and roots from 28 dpi WT, *mot1.2-1*, and *mot1.2-1* transformed with *MtMOT1.2-HA* plants. Data are the mean ± SD of at least 2 sets of 6 pooled independently transformed plants. (D) Nitrogenase activity of 28 dpi WT, *mot1.2-1*, and *mot1.2-1* transformed with *MtMOT1.2-HA* plants. Data are the mean ± SD of 2 sets of 6 pooled plants. 100 % = 0.161 nmol acetylene · h^-1^ plant^-1^. * indicates a statistically significant difference (p<0.05). (E) Mo content in shoots, roots, and nodules of WT, *mot1.2-1*, and *mot1.2-1* transformed with *MtMOT1.2-HA* plants. Data are the mean ± SD of at least 2 sets of 6 pooled independently transformed plants. * indicates a statistically significant difference (p<0.05).

## DISCUSSION

Mo is an essential oligonutrient for plants. As part of the Mo cofactor (Moco), it is used by five different proteins: i) Nitrate reductase (NR), for the reduction of nitrate to nitrite, a key step in the inorganic nitrogen assimilation process; ii) sulfite oxidase (SO), which oxidizes sulfite to sulfate producing hydrogen peroxide and thus has a role in ROS production; iii) aldehide oxidase (AO), which is related with the production of abscisic acid and auxin; iv) xanthine dehydrogenase (XDH), which catalyses hydroxylation of aldehydes and aromatic heterocycles in the purine degradation metabolic pathway; and v) the amidoxime reducing component (ARC), which catalyse the reduction of N-hydroxylated products (Mendel and Bittner, 2006; Hille et al., 2001). Consequently, plants need a regular supply of this nutrient from soil to sink organs. Mo requirements by legumes are substantially higher that those of other dicots (Clark, 1984; Tisdale et al., 1985), as a result of the synthesis of large quantities of nitrogenase by rhizobia in their root nodules (Miller et al., 1993). These bacteria use Mo to synthesize FeMoco required for nitrogenase activity (Rubio and Ludden, 2008). While much is known on how Moco and FeMoco are synthesized (Rubio and Ludden, 2008; Mendel, 2013) much less is known on how Mo is ferried to the enzymes synthesizing each cofactor. A major breakthrough has been the identification of molybdate-specific transporters (Tejada-Jimenez et al., 2013), such as those of the MOT1 family (Tejada-Jimenez et al., 2007; Tomatsu et al., 2007) and the description of the role of these proteins in molybdate uptake from soil in *Arabidopsis* and in *L. japonicus* (Tomatsu et al., 2007; Baxter et al., 2008; Duan et al., 2017). More recently, a MOT1 protein has been reported as responsible for molybdate uptake by sink organ cells in *M. truncatula* nodules, participating in symbiotic nitrogen fixation (Tejada-Jimenez et al., 2017). However, how Mo reaches this sink organ still remains obscure.

*MtMOT1.2* is a *M. truncatula* MOT1 family member that is expressed in roots and nodules (Tejada-Jimenez et al., 2017). All MOT1 members identified so far have shown Mo transport activity (Tejada-Jiménez et al., 2007; Tomatsu et al., 2007; Baxter et al., 2008; Gasber et al., 2011; Duan et al., 2017; Tejada-Jiménez et al., 2017). Yeast transport assays confirm that MtMOT1.2 is able to transport molybdate, showing kinetic parameters comparable to those of previously characterized MtMOT1.3 and LjMOT1 transporters (Duan et al., 2017; Tejada-Jimenez et al., 2017), with lower affinity and higher speed than MOT1 proteins from *Chlamydomonas reindhartii* and *A. thaliana* (Tejada-Jimenez et al., 2007; Tomatsu et al., 2007). This difference could be due to a higher local concentration of molybdate in nodules corresponding to the increased Mo demand of these organs, for which a low-affinity system would be enough, but that would need to work at higher rates. In spite of its relatively low molybdate affinity, MtMOT1.2 is a transporter specific for this anion, since the addition of up to a 4,000-fold excess of the structurally similar anion sulfate did not inhibit Mo transport.

Within roots and nodules *MtMOT1.2* expression was detected around the vessels, as indicated by promoter::*gus* fusions and immunolocalization of a HA-tagged version of the protein. More specifically, the tagged protein could be detected in endodermal cells in both nodule and root vessels. As it was the case for *A. thaliana* MOT1 (Tomatsu et al., 2007), MtMOT1.2-HA was observed in the plasma membrane and in an endodermal compartment resembling the endoplasmic reticulum. This could indicate a role in introducing molybdate into the cytosol, either from the cell exterior or from intracellular reserves. Alternatively, the endoplasmic reticulum subpopulation of MtMOT1.2 could also correspond to newly synthesized protein being ferried towards the plasma membrane. Surprisingly for a vascular transporter, no polar localization in the cell was observed. Molybdate could be introduced into the cell from the apoplast or from the vessels. As a result, in the abscense of another driving force, it would result in a futile cycle in which no net transfer of molybdate from sink to source would occur. Since this is not what happens, molybdate is being delivered from root to nodules, it might be speculated that molybdate delivery could be driven by a net mass-effect in which the molybdate pulled from the nitrogen-fixing cells, with their high molybdate uptake capability for FeMoco synthesis, would prevent a backward flux of Mo. The net transport into rhizobia-infected cells would have to be driven by transforming molybdate into different chemical species, rather than a substrate MOT1 proteins. Such a system would also ensure that should Mo not be used and accumulated in a given compartment, it would be rapidly recycled back for use elsewhere.

The localization of MtMOT1.2 in the vasculature and its function in molybdate uptake into the cell is suggestive of a role in the sink-to-source transport of this oligonutrient. Its position in the root endodermis indicates that it would be facilitating the transfer of apoplastic molybdate to the vasculature, so that molybdate would then be transferred to leaves or nodules. However, our data indicate that MtMOT1.2 does not play an essential role in molybdate transport to the leaves, since mutant plants in this gene did not have any significant growth alteration compared to wild-type plant, and, more importantly, Mo-dependent nitrate reductase activity was not affected in *mot1.2-1* when nitrate was the sole nitrogen source to these plants. In contrast, when plants relied on symbiotic nitrogen fixation for assimilable nitrogen, *mot1.2-1* plants showed a severe growth defect. This difference in growth between the two different nutritional situations could be the result of two non-incompatible possibilities: i) MtMOT1.2 is functionally substituted by another molybdate transporter in roots when mutated, and ii) MtMOT1.2 is only essential for molybdate release to the nodule. The endodermal localization in nodule vessels and the predicted direction of transport is indicative of a role in introducing the molybdate delivered by the vessels into endodermal cells. This would be the first step towards transferring molybdate to the nodule apoplast for uptake by MtMOT1.3. The plant growth defect observed in nitrogen-fixing conditions arises from the reduction of nitrogenase activity in these plants, consequence of insufficient molybdate reaching the nitrogen-fixing cells, as indicated by the lower levels of Mo in the *mot1.2-1* nodules and the restoration of the wild-type phenotype when more Mo was added to the nutrient solution. The pattern of Mo accumulation in *mot1.2-1* plants compared to their controls, with no significant changes in roots and a decrease in nodules, indicates that the defect in molybdate delivery for nitrogen fixation is occurring at the level of nodule vessels and not in loading the root vasculature with Mo. Otherwise, an accumulation of Mo in *mot1.2* roots would be expected as well as a decrease in shoots, and none was detected in either (in this case, even slightly higher levels were detected).

In summary, MtMOT1.2 would position itself between molybdate root uptake transporter, likely MtMOT1.1 as the closest LjMOT1 orthologue, and the nodule apoplast molybdate uptake protein MtMOT1.3 (Fig. 6). MtMOT1.2 would facilitate the transfer of this oligonutrient into endodermal cells mediating the sink-to-source molybdate trafficking, which would be controlled by mass-effects to ensure that it reaches its destination. However, a critical point remains to be solved, which is the identity of the proteins mediating molybdate efflux from the cytosol to the symbiosome. Whether these are sulfate transporters, or whether a novel family of Mo transporters with a direction of transport opposite to MOT1 proteins, remains to be unveiled.

**Figure 6.**
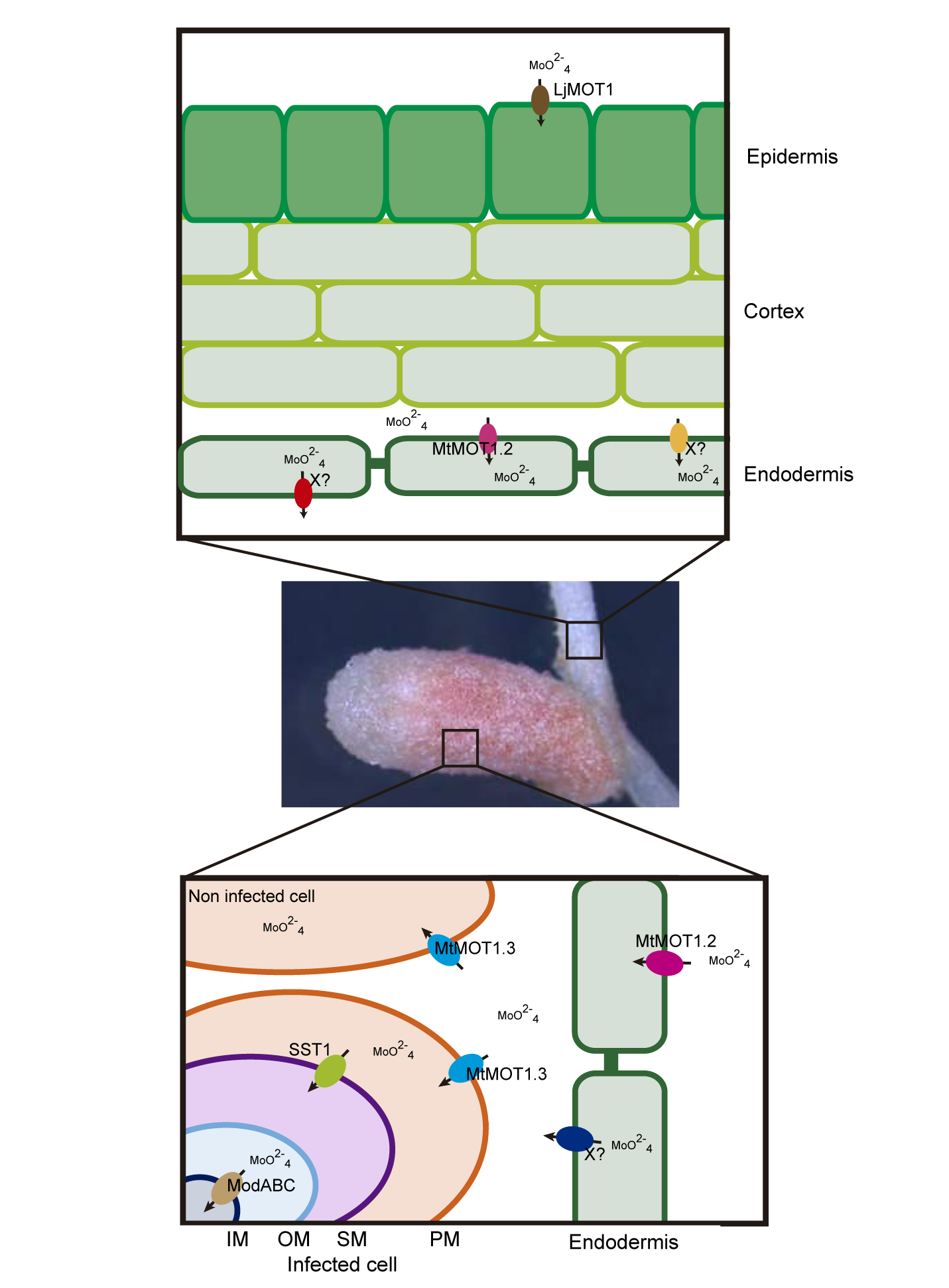
Model of molybdate transport in *Medicago truncatula*. Upper panel, molybdate uptake from soil would be mediated by epidermal root transporters similar to LjMOT1, perhaps MtMOT1.1. Symplastically and apoplastically-delivered molybdate will reach the endodermis were MtMOT1.2 will introduce it from the apoplast. This role is not carried out only by this protein, another molybdate transporter would also participate delivering molybdate to the shoots. Molybdate efflux from the endodermis into the xylem is mediated by a yet-to-be determined transporter. Lower panel, once molybdate reaches the nodule, it is recovered from the vasculature by MtMOT1.2 that would introduce it into the endodermal cells. Through an unknown protein, this molybdate is released into the apoplast, where MtMOT1.3 will introduce it into nodule cells. SST1 and bacterial ModABC would then direct the cytosolic molybdate to nitrogen-fixing bacteroids.

## METHODS

### Biological material and growth conditions

*M. truncatula* R108 seeds were scarified by incubating with concentrated sulfuric acid for 7 min. After several washes with cold water, the seed surfaces were sterilized in 50 % (v/v) bleach for 90 s, and left in sterile water in the dark overnight, followed by a 48 h incubation at 4 °C. Seed germination was done in water-agar plates 0.8 % (w/v). Seedlings were planted in sterile perlite pots, and inoculated with *Sinorhizobium meliloti* 2011 or the same bacterial strain transformed with pHC60 (Cheng and Walker, 1998). Plants were grown in a greenhouse with 16 h of light and 22 °C, and watered every two days with Jenner’s solution or water alternatively (Brito et al., 1994). Nodule collection was carried out at 28 dpi. Non-nodulated plants were supplemented every two weeks with 2 mM KNO_3_, instead of being inoculated with *S. meliloti* 2011. For hairy root transformation of *M. truncatula* seedlings, *Agrobacterium rhizogenes* strain ARqua1 having the appropriate vector was used (Boisson-Dernier et al., 2001). Agroinfiltration experiments for transitory expression were done in *N. benthamiana* leaves using *A. tumefaciens* C58C1 as a vector for the corresponding genetic construct. *N. benthamiana* plants were grown in the greenhouse under the same conditions as *M. truncatula.*

In heterologous expression assays the yeast *S. cerevisiae* strain 31019b (MATa *ura3 mep1Δ mep2Δ::LEU2 mep3Δ:: KanMX2*) was used (Marini et al., 1997). Yeasts were grown in synthetic dextrose (SD) or yeast peptone dextrose (YPD) medium supplemented with 2 % glucose (Sherman et al., 1986).

### Molybdate uptake

*S. cerevisiae* cells grown in SD medium were transferred to 10 mM MES-Ca(OH)_2_ buffer (pH 5.8) containing 0.1 mM MgCl_2_, 2 mM CaCl_2_ and 0.5 % glucose. Molybdate uptake was measured after 30 min incubation at 28 °C in the presence of 500 nM Na_2_MoO_4_. For molybdate transport kinetics yeast cells were transferred to the same MES buffer supplemented with 100, 200, 500, 1000 and 2000 nM Na_2_MoO_4_ and incubated for 30 min at 28 °C. Molybdenum determination was carried out in 10 mL cell-free MES buffer using the method previously described (Cardenas and Morteson, 1975)

### GUS Staining

A transcriptional fusion between *MtMOT1.2* promoter region and the *gus* gene was obtained by amplifying 1.4 kb upstream of the *MtMOT1.2* start codon using the primers 5MtMOT1.2-1446GW and 3MtMOT1.2pGW (Supplemental Table S1). This amplicon was inserted into pGWB3 (Nakagawa et al., 2007) using the Gateway cloning technology (Invitrogen). Roots were transformed as indicated above. Visualization of GUS activity was done in 28 dpi plants as described (Vernoud et al., 1999). Clearing of nodule sections was carried out with a 50 % bleach treatment for 30 min.

### Immunohistochemistry and confocal microscopy

Plasmid pGWB13 (Nakagawa et al., 2007) was used to clone a DNA fragment containing *MtMOT1.2* full gene and 1,446 kb upstream of its start codon using Gateway cloning technology (Invitrogen), adding three C-terminal hematoagglutinin (HA) epitopes in frame to MtMOT1.2. *M. truncatula* root transformation was carried out as indicated above. Transformed plants were inoculated with *S. meliloti* 2011 containing plasmid pHC60 that constitutively expresses GFP. Nodules and roots, collected at 28 dpi, were fixed in 4 % (w/v) paraformaldehyde and 2.5 % (w/v) sucrose in phosphate-buffered saline (PBS) at 4 °C and left overnight. Fixed plant material was sectioned with a Vibratome 1000 Plus (Vibratome), in 100 µm sections. Dehydration of the sections was performed by incubation with methanol dilution series (30 %, 50 %, 70 % and 100 % in PBS) for 5 min and then rehydrated following the same methanol series in reverse order. Cell wall permeabilization was done by incubating with 2 % (w/v) cellulase in PBS for 1 h, and 0.1 % (v/v) Tween 20 in PBS for 15 min. Bovine serum albumin 5% (w/v) was used to block the sections. As primary antibody, a 1:50 dilution in PBS of anti-HA mouse monoclonal antibody (Sigma) was used. This dilution was incubated with the sections for 2 h at room temperature and then washed away three times with PBS for 10 min. Secondary antibody used was 1:40 Alexa594-conjugated anti-mouse rabbit monoclonal antibody (Sigma) in PBS. The incubation was performed at room temperature for 1 h and then sections were washed three times with PBS for 10 min. DNA was stained using DAPI. Images were obtained with a confocal laser-scanning microscope (Leica SP8).

### Gold-immunohistochemistry and electron microscopy

Plants were transformed with plasmid pGWB13 containing *MtMOT1.2* full gene and 1,446 bp upstream of its start codon. Transformed plants were inoculated with *S. meliloti* 2011. Nodules were collected at 28 dpi and were fixed in 1 % formaldehyde and 0.5 % glutaraldehyde in 50 mM potassium phosphate (pH 7.4) for 2 h. After that the fixation solution was renewed for 1.5 h. Samples were washed in 0.05 M potassium phosphate (pH 7.4) 3 times during 30 min and 3 times for 10 min. Nodules were dehydrated by incubation with ethanol dilution series of 30 %, 50 %, 70 %, 90 % during 10 min, 96 % for 30 min and 100 % during 1 h. Samples were included in a series of ethanol and LR–white resin (London Resin Company Ltd, UK) dilutions: 1:3 during 3 h, 1:1 were left overnight and 3:1 during 3 h. Nodules were included in resin during 48 h. All the process was performed at 4 °C. Nodules were placed in gelatine capsules and filled with resin and polymerized at 60 °C for 24 h. One-micron thin sections were cut at Centro Nacional de Microscopia Electrónica (Spain) with Reichert Ultracut S-ultramicrotome fitted with a diamond knife. Thin sections were blocked in 2 % bovine serum albumin in phosphate buffer saline (PBS) for 30 min. As primary antibody, a 1:20 dilution in PBS of anti-HA rabbit monoclonal antibody (Sigma) was used. Samples were washed 10 times in PBS for 2 min. Secondary antibody used was 1:150 anti-rabbit goat conjugated to a 15 nm gold particle (BBI solutions) diluted in PBS. Incubation was performed for 1 h, after that samples were washed 10 times in PBS for 2 min and 15 times in water for 2 min. Sections were stained with 2 % uranyl acetate and visualised in a JEM 1400 electron microscope at 80 kV.

### Transient expression in *Nicotiana benthamiana* leaves

Experiment was performed as is described by Wood et al (2009). GFP was fused to the C terminus of *MtMOT1.2* coding sequence by cloning it into pGWB5 (Nakagawa et al., 2007) by Gateway cloning technology (Invitrogen). Four-week-old *N. benthamiana* leaves were injected with *A. tumefaciens* C58C1 (Deblaere et al., 1985) cells independently transformed with MtMOT1.2-GFP, with the plasma membrane marker pm-CFP pBIN (Nelson et al., 2007) or with the silencing suppressor p19 of *Tomato bushy stunt virus* (Wood et al., 2009). Expression in the leaves was analyzed after 3 d by confocal laser-scanning microscopy (Leica SP8).

### Nitrogenase activity

Nitrogenase activity was measured by the acetylene reduction assay (Hardy et al., 1968). 28 dpi wild-type and mutant plants were introduced in 30 ml tubes and sealed with rubber stoppers. Each tube contained at least four independently transformed plants. 10 % of the gas phase from each bottle was replaced by the same volume of acetylene. Tubes were incubated for 30 min at room temperature. Ethylene production was measured by analyzing 0.5 ml gas samples with a Shimadzu GC-8A gas chromatograph using a Porapak N column (Shimadzu, Kyoto, Japan).

### Metal content determination

Metal content was determined by inductively coupled plasma optical emission spectrometry in three sets of 28 dpi roots, shoots, and nodules, each set originating from a pool of five plants. The experiment was carried out at the Unit of Metal Analysis in the Scientific and Technology Centers of the Universidad de Barcelona (Spain). These samples were digested with HNO_3_, H_2_O_2_ and HF in a Teflon reactor at 90 °C. The sample was diluted with deionized water. Final volume was calculated by weight and weight: volume ratios. The samples were digested with three blanks in parallel. Metal determination was carried out in an Agilent 7500cw instrument under standard conditions. Calibration was carried out with five solutions prepared from certified NIST standards.

### Nitrate reductase activity

Nitrate reductase activity was analyzed as described by Tejada-Jimenez *et al* (2017). Briefly, a crude extract was obtained from approximately 100 mg of fresh material in 100 mM potassium phosphate, pH 7.5, 5 mM magnesium acetate, 10% glycerol (v/v), 10 % polyvinylpolypyrrolidone (w/v), 0.1% Triton X-100, 1 mM EDTA,0.05 % ß-mercaptoethanol and 1 mM PMSF. Plant material was homogenized with liquid nitrogen and 1:6 extraction buffer (v/v), and centrifuged at 14,000 x*g* at 4 °C for 15 min. The reaction was started adding 50 μl of crude extract to 0.5 ml of reaction buffer and incubated at 30 °C for 20 min. The reaction buffer contained 50 mM potassium phosphate, pH 7.5, 10mM KNO_3_, 5 mM EDTA and 0.5 mM NADH. The reduction reaction was stopped by adding 1 volume of 1 % sulfanilamide in 2.4 M HCl, and 1 volume of 0.02 % N-1-naphtyl-ethylenediamine. After centrifugation, the supernatant was collected and its absorbance at 540 nm measured in a UV/visible spectrophotometer (Ultrospect 3300 pro; Amersham Bioscience).

### Statistical analysis

Student’s unpaired *t*-test was used to calculate statistical significance of observed differences. Test results with p-values less than 0.05 were considered as statistically significant.

## ACKNOWLEDGEMENTS

The authors would like to thank Dr. Emilio Fernández and Dr. Aurora Galván (Universidad de Córdoba) for their help with the yeast transport assays, as well as to members of Laboratory 281 at Centro de Biotecnología y Genómica de Plantas (UPM-INIA) for their support and feed-back in preparing this manuscript.

## References

Baxter I, Muthukumar B, Park HC, Buchner P, Lahner B, Danku J, Zhao K, Lee J, Hawkesford MJ, Guerinot ML, Salt DE (2008) Variation in molybdenum content across broadly distributed populations of *Arabidopsis thaliana* is controlled by a mitochondrial molybdenum transporter (MOT1). PLoS Genet. 4 (2): e1000004.

Bernard SM, Habash DZ (2009) The importance of cytosolic glutamine synthetase in nitrogen assimilation and recycling. New Phytol. 182: 608–620.

Boisson-Dernier A, Chabaud M, Garcia F, Becard G, Rosenberg C, Barker DG (2001) *Agrobacterium rhizogenes*-transformed roots of *Medicago truncatula* for the study of nitrogen-fixing and endomycorrhizal symbiotic associations. Mol Plant Microbe Interact 14: 695–700.

Brear EM, Day DA, Smith PM (2013) Iron: an essential micronutrient for the legume-rhizobium symbiosis. Front Plant Sci 4: 359

Brito B, Palacios JM, Hidalgo E, Imperial J, Ruiz-Argueso T (1994) Nickel availability to pea (*Pisum sativum* L.) plants limits hydrogenase activity of *Rhizobium leguminosarum* bv. viciae bacteroids by affecting the processing of the hydrogenase structural subunits. J Bacteriol 176: 5297–5303.

Cardenas J, Mortenson LE (1975) Role of molybdenum in dinitrogen fixation by Clostridium pasteurianum. J Bacteriol 123 (3): 978–984.

Cheng HP, Walker GC (1998) Succinoglycan is required for initiation and elongation of infection threads during nodulation of alfalfa by Rhizobium meliloti. J Bacteriol 180: 5183–5191.

Cheng X, Wen J, Tadege M, Ratet P, Mysore KS (2011) Reverse genetics in *Medicago truncatula using* Tnt1 insertion mutants. Methods Mol Biol. 678: 179–190.

Cheng X, Wang M, Lee HK, Tadege M, Ratet P, Udvardi M, Mysore KS, Wen J (2014) An efficient reverse genetics platform in the model legume Medicago truncatula. New Phytol. 201: 1065–1076.

Clark, R.B (1984) Physiological aspects of calcium, magnesium and molybdenum deficiencies in plants. Soil Acidity and Liming 12: 99–170.

Deblaere R, Bytebier B, De Greve H, Deboeck F, Schell J, Van Montagu M, Leemans J (1985) Efficient octopine Ti plasmid-derived vectors for Agrobacterium-mediated gene transfer to plants. Nucleic Acids Res 13: 4777–4788.

Downie JA (2014) Legume nodulation. Curr. Biol. 24: R184–R190.

Duan G, Hakoyama T, Kamiya T, Miwa H, Lombardo F, Sato S, Tabata S, Chen Z, Watanabe T, Shinano T, Fujiwara T (2017) LjMOT1, a high-affinity molybdate transporter from Lotus japonicus, is essential for molybdate uptake, but not for the delivery to nodules. Plant J 90: 1108–1119.

Gao JS, Wu FF, Shen SL, Meng Y, Cai YP, Lin Y (2016) A putative molybdate transporter LjMOT1 is required for molybdenum transport in Lotus japonicus. Physiol. Plant. 158: 331–340.

Gasber A, Klaumann S, Trentmann O, Trampczynska A, Clemens S, Schneider S, Sauer N, Feifer I, Bittner F, Mendel RR (2011) Identification of an Arabidopsis solute carrier critical for intracellular transport and inter-organ allocation of molybdate. Plant Biol 13: 710–718.

González-Guerrero M, Matthiadis A, Sáez A, Long TA (2014) Fixating on metals: new insights into the role of metal in nodulation and symbiotics nitrogen fixation. Front Plant Sci 5: 45.

Hardy RW, Holsten RD, Jackson EK, Burns RC (1968) The acetylene-ethylene assay for n(2) fixation: laboratory and field evaluation. Plant Physiol 43: 1185–1207.

Hille R, Nishino T, Bittner F (2011) Molybdenum enzymes in higher organisms. Coord Chem Rev 225: 1179–1205.

Kaiser BN, Gridley KL, Ngaire BJ. Phillips T, Tyeman SD (2005) The role of molybdenum in agricultural plant production. Ann Bot 96: 754–54.

Limpens E, Ivanov S, van Esse W, Voets G, Fedorova E. Bisseling T (2009) Medicago N_2_-Fixing symbiosomes acquire the endocytic identity marker Rab7 but delay the acquisition of vacuolar identity. Plant Cell 21: 2811–2828.

Marini AM, Soussi-Boudekou S, Vissers S, Andre B (1997) A family of ammonium transporters in Saccharomyces cerevisiae. Mol. Cell. Biol 17: 4282–4293.

Mendel RR, Hansch R (2002) Molybdoenzymes and molybdenum cofactor in plants. J Exp Bot 53: 1689–1698.

Mendel RR, Bittner F (2006) Cell biology of molybdenum. Biochim Biophys Acta 1763: 621–635.

Mendel RR (2013) The molybdenum cofactor. J Biol Chem 288 (19): 13165–13172.

Miller RW, Yu Z, Zarkadas CG (1993) The nitrogenase proteins of Rhizobium meliloti: purification and properties of the MoFe and Fe components. Biochim Biophys Acta 1163: 31–41.

Nakagawa T, Kurose T, Hino T, Tanaka K, Kawamukai M, Niwa Y, Toyooka K, Matsuoka K, Jinbo T, Kimura T (2007) Development of series of gateway binary vectors, pGWBs, for realizing efficient construction of fusion genes for plant transformation. J Biosci Bioeng 104: 34–41.

Nelson BK, Cai X, Nebenfuhr A (2007) A multicolored set of in vivo organelle markers for co-localization studies in Arabidopsis and other plants. Plant J 51: 1126–1136.

O’Hara GW (2001) Nutritional constraints on root nodule bacteria affecting symbiotic nitrogen fixation: a review. Aust. J. Exp. Agric. 41: 417–433

Oldroyd GED (2013) Speak, friend, and enter: signalling systems that promote beneficial symbiotic associations in plants. Nat Rev Micro 11: 252–263.

Roth LE, Stacey G (1989) Bacterium release into host cells of nitrogen-fixing soybean nodules: the symbiosome membrane comes from three sources. Eur J Cell Biol 49: 13–23

Roux B, Rodde N, Jardinaud MF, Timmers T, Sauviac L, Cottret L, Carrere S, Sallet E, Courcelle E, Moreau S (2014) An integrated analysis of plant and bacterial gene expression in symbiotic root nodules using laser-capture microdissection coupled to RNA sequencing. Plant J 77: 817–837.

Rubio LM, Ludden PW (2005) Maturation of nitrogenase: a biochemical puzzle. J. Bacteriol. 187: 405–414.

Rubio LM, Ludden PW (2008) Biosynthesis of the iron-molybdenum cofactor of nitrogenase. Annu. Rev. Microbiol. 62: 93–111.

Sherman F, Fink GR, Hicks JB (1986) Methods in yeast genetics. Plainview, NY: Cold Spring Harbor Lab Press.

Stiefel EI (2002) Molybdenum and tungsten: their roles in biological processes. In: Sigel A, Sigel H, eds. Metal ions in biological systems New York: Marcel Dekker Inc., 1–30.

Stout PR, Meagher WR, Pearson GA, Johnson CM (1951) Molybdenum nutrition of crop plants: I. The influence of phosphate and sulfate on the absortion of molybdenum from soils and solution cultures. Plant Soil 3: 51–87.

Tadege M, Wen J, He J, Tu H, Kwak Y, Eschstruth A, Endre G, Zhao PX, Chabaud M, Ratet P, Mysore KS (2008) Large scale insertional mutagenesis using the Tnt1 retrotransposon in the model legume Medicago truncatula. Plant Journal 54: 335–347.

Tejada-Jimenez M, Llamas A, Sanz-Luque, E, Galvan A, Fernandez E (2007) A high-affinity molybdate transporter in eukaryotes. Proc Natl Acad Sci U S A 104: 20126–20130.

Tejada-Jiménez M, Galván A, Fernández E (2011) Algae and humans share a molybdate transporter. Proc Natl Acad Sci U S A 108: 6420–6425.

Tejada-Jimenez M, Chamizo-Ampudia A, Galvan, A, Fernandez E, Llamas A (2013) Molybdenum metabolism in plants. Metallomics 5, 1191–1203

Tejada-Jimenez M, Gil-Diez P, León-Mediavilla J, Wen J, Mysore KS, Imperial J, González-Guerrero M (2017) Medicago truncatula MOT1.3 is a plasma membrane molybdenum transporter required for nitrogenase activity in root nodules under molybdenum deficiency. New Phytol 216: 1223–1235.

Tisdale SL, Nelson WL, Beaton JD (1985) Soils fertility and fertilizers. New York, Macmillan, 4th edition: 188-239.

Tomatsu H, Takano J, Takahashi H, Watanabe-Takahashi A, Shibagaki N, Fujiwara T. (2007) An Arabidopsis thaliana high-affinity molybdate transporter required for efficient uptake of molybdate from soil. Proc Natl Acad Sci U S A 104: 18807–18812.

Udvardi M, Poole PS (2013) Transport and metabolism in legume-rhizobia symbioses. Annu. Rev Plant Biol 64: 781–805.

Vasse J, de Billy F, Camut S, Truchet G (1990) Correlation between ultrastructural differentiation of bacteroids and nitrogen fixation in alfalfa nodules. J Bacteriol 172: 4295–4306.

Vernoud V, Journet EP, Barker DG (1999) MtENOD20, a Nod factor-inducible molecular marker for root cortical cell activation. Mol Plant Microbe Interact 12: 604–614.

Wood CC, Petrie JR, Shrestha P, Mansour MP, Nichols PD, Green AG, Singh SP (2009). A leaf-based assay using interchangeable design principles to rapidly assemble multistep recombinant pathways. Plant Biotechnol J 7: 914–924.

